# Trend, Population Structure and Trait Mapping from 15 Years of National Varietal Trials of UK Winter Wheat

**DOI:** 10.1101/2021.05.17.444481

**Authors:** Oluwaseyi Shorinola, James Simmonds, Luzie U. Wingen, Keith Gardner, Cristobal Uauy

**Author notes:** **Corresponding Author**: Oluwaseyi Shorinola, John Innes Centre, Norwich Research Park, Norwich UK, NR4 7UH,.

## Abstract

There are now a rich variety of genomic and genotypic resources available to wheat researchers and breeders. However, the generation of high-quality and field-relevant phenotyping data which is required to capture the complexities of gene x environment interactions remains a major bottleneck. Historical datasets from national variety performance trials (NVPT) provide sufficient dimensions, in terms of numbers of years and locations, to examine phenotypic trends and study gene x environment interactions. Using NVPT for winter wheat varieties grown in the UK between 2002 – 2017, we examined temporal trends for eight traits related to yield, adaptation, and grain quality performance. We show a non-stationary linear trend for yield, grain protein content, HFN and days to ripening. Our data also show high environmental stability for yield, grain protein content and specific weight in UK winter wheat varieties and high environmental sensitivity for Hagberg Falling Number. Using the historical NVPT data in a genome-wide association analysis, we uncovered a significant marker-trait association peak on wheat chromosome 6A spanning the *NAM-A1* gene that have been previously associated with early senescence. Together our results show the value of utilizing the data routinely collected during variety evaluation process for examining breeding progress and the genetic architecture of important traits.

## INTRODUCTION

Over the last three years, there has been a rapid surge in the development of genomic resources for wheat (reviewed in Adamski *et al.* 2020). This includes a chromosome-scale reference assembly of the Chinese Spring cultivar (RefSeqv1) and a pan-genome resource comprised of chromosome and scaffold-level assemblies of 15 hexaploid wheat cultivars (IWGSC *et al.* 2018; Walkowiak *et al.* 2020). There is also a wide range of array-based (Axiom-35K, iSelect 90K, Axiom-660K and Axiom-820K; Wang *et al.* 2014; Winfield *et al.* 2016; Allen *et al.* 2017), sequencing-based (e.g DARTSeq, RADSeq) or PCR-based (e.g KASP, TaqMan, rhAmp; Semagn *et al.* 2014; Ayalew *et al.* 2019) SNP genotyping assays available to wheat researchers and breeders. There have also been efforts to re-sequence different wheat populations either through reduced-representation sequencing approach like exome-capture and sequencing (e.g He *et al.* 2019; Krasileva *et al.* 2017; Jordan *et al.* 2015) or through whole genome resequencing (e.g Cheng *et al.* 2019; Scott *et al.* 2020). This preponderance of genomics and genotypic data which are available in open-access repositories (e.g. EnsemblPlants, CerealsDB; Bolser *et al.* 2016; Howe *et al.* 2020; Wilkinson *et al.* 2020) now makes it possible to map traits at high-resolution (e.g Walkowiak *et al.* 2020), examine population diversity at whole genome levels or in breeding units (haplotypes: e.g Brinton *et al.* 2020; Scott *et al.* 2020), and implement genome-assisted breeding schemes using marker-assisted and/or genomic selection (e.g Sweeney *et al.* 2019; Rasheed and Xia 2019).

Despite these advances, the generation of high-quality and field-relevant phenotyping data remains a major bottleneck. Modern phenomics platforms have improved phenotyping throughput and precision under controlled conditions, but these do not always capture the environmental effects experienced under real-world farming conditions (Yang *et al.* 2020). Given climate change projections of fluctuating radiation, heat and precipitation patterns in major wheat growing areas (including the UK), breeding for phenotypic stability and understanding complex gene x environment interactions is of high priority (Semenov 2009; Trnka *et al.* 2019).

Due to their large scale and multi-environment (years and locations) design, historical dataset from national variety performance trials (NVPT) provide sufficient dimensions, in terms of years and locations to examine phenotypic trends and study gene x environment interactions. These historical datasets are, however, incomplete by design because of, for example, changes in the number and specific set of varieties trialed and changes in the field sites used from year to year. Previous studies have analyzed NPVT data for wheat in the UK (Silvey 1981; Mackay *et al.* 2011) and similar analyses of historical data have been conducted elsewhere (e.g Crossa *et al.* 2007; Pozniak *et al.* 2012).

In the UK, new wheat varieties undergo statutory tests before they are registered on the National List (NL). Registered varieties are subsequently introduced (or maintained on) the UK Recommended List (RL) after undergoing independent non-statutory NPVT managed by the Agriculture and Horticulture Development Board (AHDB, formerly Home-Grown Cereals Authority). The NL serves as variety registry while the RL is used as a reference by farmers for variety selection. Mackay et al. (2011) re-analyzed data from the UK NL and RL trials conducted between 1948 – 2007, and found significant yield improvement that was mostly attributed to plant breeding. In the present study, we analyzed data from the UK RL NVPT for winter wheat between 2002 - 2017 and used this to examine temporal trends in eight yield, adaptation, and grain quality traits. We also demonstrate the usefulness of these NVPT dataset for trait mapping to uncover loci of breeding importance.

## MATERIALS AND METHODS

### NVPT datasets

We downloaded result files for the NVPT for winter wheat in the UK from 2002 – 2010 and 2012 - 2017 from the AHDB website (accessible at: https://ahdb.org.uk/knowledge-library/recommended-lists-for-cereals-and-oilseeds-rl-harvest-results-archive). We focused our study on data for eight traits including yield, adaptation and grain quality traits. Yield and height data were collected from treated and untreated trials. The treated trials included management for diseases (fungicide spray) while the untreated trials did not include disease management. Both trials were managed under standard husbandry practices including the application of plant growth regulator (PGR), herbicide, fertiliser and pest control management as recommended by AHDB. Details of the AHDB RL trial protocol is accessible at: https://ahdb.org.uk/rlprotocols. Before analyses, we filtered the dataset to remove observations with unknown locations or from locations where trials were abandoned. Varieties that were trialed in a single year were also omitted. The nomenclature of varieties, locations and counties were standardized in cases where different designation or acronyms were used for the same variety, location or county across different years. After filtering, the distribution of the observations obtained for each of the eight target traits resemble a bell curve suggesting normal distribution (Figure S1).

### Germplasm

Data for a total of 168 varieties were used in this study. These include 133 varieties whose phenotype information were obtained from the AHDB website as described above. For 139 varieties, which included additional 35 pre-2002 UK wheat varieties, genotype data from the Axiom-35K array was used as described below. The number of varieties used for each analysis in this study are detailed in Figure S2.

### Statistical Analyses

We used a two-stage approach to examine the linear trend of trait from the NVPT data. First, we fitted a linear mixed model (LMM) to the NVPT data using restricted maximum likelihood (REML) estimation. The model was implemented using the lme4 package in R as:

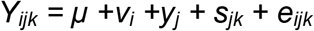

*Y_ijk_* is the historical performance of variety *i* in year *j* at location *k*. *μ* is the overall mean performance of all varieties, *v_i_* is the effect of variety *i*, y_*j*_ is the effect of year *j* (the calendar year of the trial) and *S_jk_* is the effect of location *k* within year *j*. e_*ijk*_ is the residual variance arising from factors not accounted for in the model including variety x year interaction. As our main interest was the performance for each variety, the variety effect was fitted as fixed factor while the year and site (nested within year) were fitted as random factors. This is slightly different to the strategy used by Mackay et al. (2011), which also included calendar year as a fixed factor to account for the long year interval (1948 – 2002) examined and changes in trial management system across these years. Given the short interval examined in this study, we believe the management systems were fairly uniform across the trial year. We derived estimates for the varieties means (EVM) from the LMM. Second, we used a linear model to regress the EVM derived from the LMM above against the year the variety was first entered into the NVPT. For trait comparison between end-use groups, Analysis of Variance (ANOVA) followed by post-hoc TukeyHSD was used to evaluate and compare significant difference in EVM of varieties belonging to different end-use groups. The lstrend function implemented in the *R* lsmeans package (Lenth 2016) was used to estimate and compare slopes of the linear regression between groups. For slope comparisons between the four end-use groups, the adjusted P value is presented based on Tukey’s method of comparison.

We used the Finlay Wilkinson regression to examine phenotype stability (Finlay and Wilkinson 1963). The original Finlay Wilkinson regression used by breeders to examine varietal adaptability is not best suited for data from incomplete trial design as the environment means used for normalizing varietal performance are biased due to incomplete replication of varieties across all environments. To circumvent this bias in our analysis, we used the Bayesian method proposed by Su et al., (2006) and implemented in the *R* package FW (Lian and de los Campos 2016). Only varieties that were trialed in more than three years were used for this analysis. The mean values for each variety in each year were used as input. The model was fitted with the Bayesian “gibbs” method, with 50000 iterations and 5000 burnIn rate as suggested for wheat analyses in the FW package paper (Lian and de los Campos 2016). The FW coefficients are presented as b + 1 which describes expected change in variety performance per unit change of the environment effect (Lian and de los Campos 2016).

### Genotyping, population structure and association analysis

A subset of 139 modern varieties and historic cultivars were genotyped using the Axiom-35K array (Allen *et al.* 2017). We filtered the genotype data to include only sites with > 0.05 minor allele frequencies. Marker with heterozygous calls but that were missing one of the homozygous calls (e.g markers with AA and AC but missing CC) were also removed as these are likely due to wrong genotype assignment during automated genotype cluster analysis. The markers were filtered to remove pair of loci with high linkage disequilibrium (*R*^2^ > 0.75). This was done to remove biases arising from high LD loci (such as from introgression from wild relatives) that can bias the contributions of such loci in population structure analysis. To assign physical positions to the Axiom markers, their sequences were used as queries in BLASTn alignments against the IWGSC RefSeqv1.0 assembly (IWGSC *et al.* 2018) as described in Brinton et al. (2020) and the best hits on each of the three wheat homoeologous genomes (A, B and D) were recorded. Of these, the correct homoeologous chromosome was selected using genetic mapping information from 13 populations (Gardiner *et al.* 2019) where available for each marker. Otherwise, the highest BLASTn score was used to select the homoeologous chromosome. In case of conflicting genetic mapping results for the correct chromosome between the mapping populations, the most frequent outcome was used.

Population structure analysis was done using discriminant analysis of principal component (DAPC) as implemented in the Adegenet *R* package (Jombart and Ahmed 2011). For this, the number of population cluster (*k*) was determined by kmeans clustering using a range of *k*. The *k* with the minimum Bayesian Information Criterion was selected as the optimum *k*. To increase the accuracy of grouping, 50 iterations of the kmeans clustering algorithm was run and the population group to which a variety was most frequently assigned was selected. Also, the cross-validation function (xvalDapc) was used to select the optimum number of principal components to use for DAPC.

GWASpoly – a *R* package for association analysis in polyploid crop, was used for GWAS (Rosyara *et al.* 2016). We used a K+Q mixed model where K represents the kinship matrix describing the relatedness between the varieties and Q represents the population grouping derived from the DAPC analysis. A Bonferroni threshold with adjusted *P* value below 0.05 was used to select markers with significant association with the trait of interest.

### Data Availability

The original data files for the trials described in this study can be downloaded from the AHDB website at: https://ahdb.org.uk/knowledge-library/recommended-lists-for-cereals-and-oilseeds-rl-harvest-results-archive. As data for different traits are combined in these original files, we re-organized the files to separate the data for each trait into separate files. The re-organized files are available at Zenodo: https://doi.org/10.5281/zenodo.4761528. The QC-filtered trial data used for subsequent analyses are presented in Table S1. Table S2 contains the end-use group information and linear-mixed-model-derived EVM for the varieties trialed. Table S3 contains the FW coefficients for each variety used in the FW regression analysis. The filtered Axiom-35K genotyping data and their genome distribution are presented in Table S4 and Table S5, respectively. Table S6 contains the population group information for each variety genotyped.

## RESULTS

### Estimates from multi-environment trial capture expected relationship between traits

We analyzed the historical data set of the UK RL NVPT from 2002 to 2017. We focused our analyses on six traits of agronomic and economic importance: yield, plant height, days to ripening, Hagberg Falling Number (HFN), grain protein content and specific weight. For yield and plant height, we analyzed data coming from (fungicide) treated and untreated trials. This results in a final dataset for eight traits. After quality controls (described in Materials and Methods), we retained 52,152 observations for these eight traits from 133 winter wheat varieties (Table S1). These 133 varieties were phenotyped in at least two years across a combined 162 locations, with a subset of 95 locations being used for evaluations in two or more years. Table 1 details the number of varieties phenotyped for each trait and the number of locations and year-location combinations used. The trial locations were spread across 43 counties and unitary authorities in England, Wales, Scotland, and Northern Ireland as shown in Figure 1.

**TABLE 1:** NUMBER OF VARIETIES, SITES, AND YEARS OF TRIALS FOR THE UK NVPT BETWEEN 2002-2017

| Trait | Varieties<sup>*</sup> | Trial Locations<sup>++</sup> | Trial Years | Year x Location Combinations | Total Observations |
| --- | --- | --- | --- | --- | --- |
| Treated yield | 133 | 158 | 15 | 410 | 13080 |
| Untreated yield | 131 | 53 | 15 | 124 | 4156 |
| Protein content | 128 | 99 | 15 | 230 | 7142 |
| Days to ripening | 133 | 108 | 15 | 247 | 7977 |
| HFN | 128 | 99 | 15 | 227 | 7154 |
| Specific weight | 129 | 99 | 15 | 231 | 7091 |
| Treated height | 108 | 74 | 11 | 171 | 3905 |
| Untreated height | 107 | 30 | 11 | 75 | 1647 |

<sup>\*</sup>Not all varieties were tested for each trait, and in each year and location.
<sup>++</sup>Some locations were used in more than one year.

**Figure 1:**
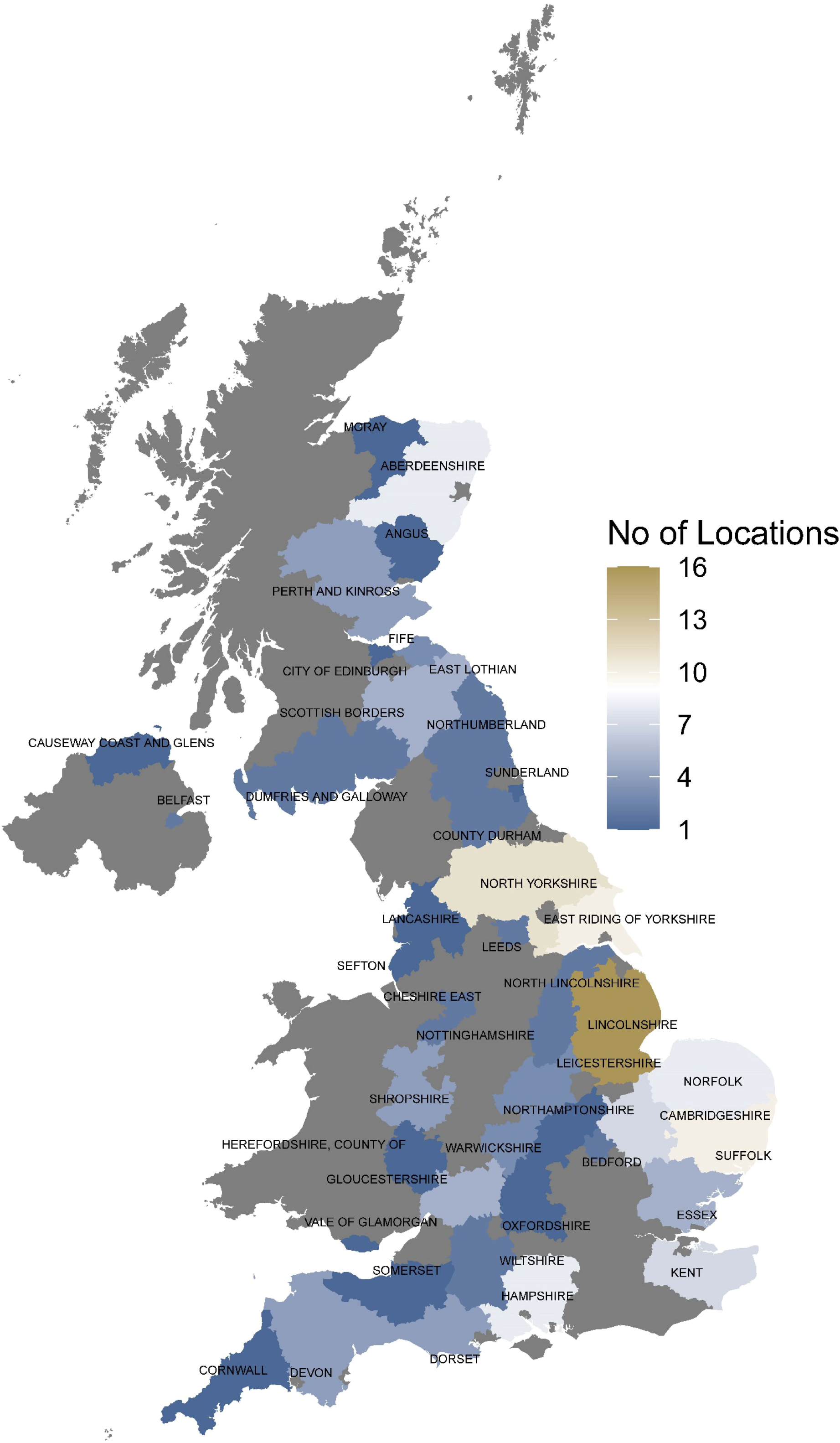
Distribution of 162 NVPT locations used in this study (2002 and 2017). The number of field sites within each county and unitary authority are indicated in colour.

Using a linear mixed model that accounted for variation arising from the different years and trial locations, we derived estimates for variety mean (hereafter referred to as EVM) for each variety for each trait (Table S2). Correlation analysis using the EVM captured expected patterns of relationship between the measured traits (Figure 2). We observed significant positive correlations between treated and untreated trials for height and yield, although the correlation between treated/untreated trials for height was much stronger than for yield. HFN and grain protein content were positively correlated to each other, but negatively correlated to treated yield, treated plant height and days to ripening.

**Figure 2:**
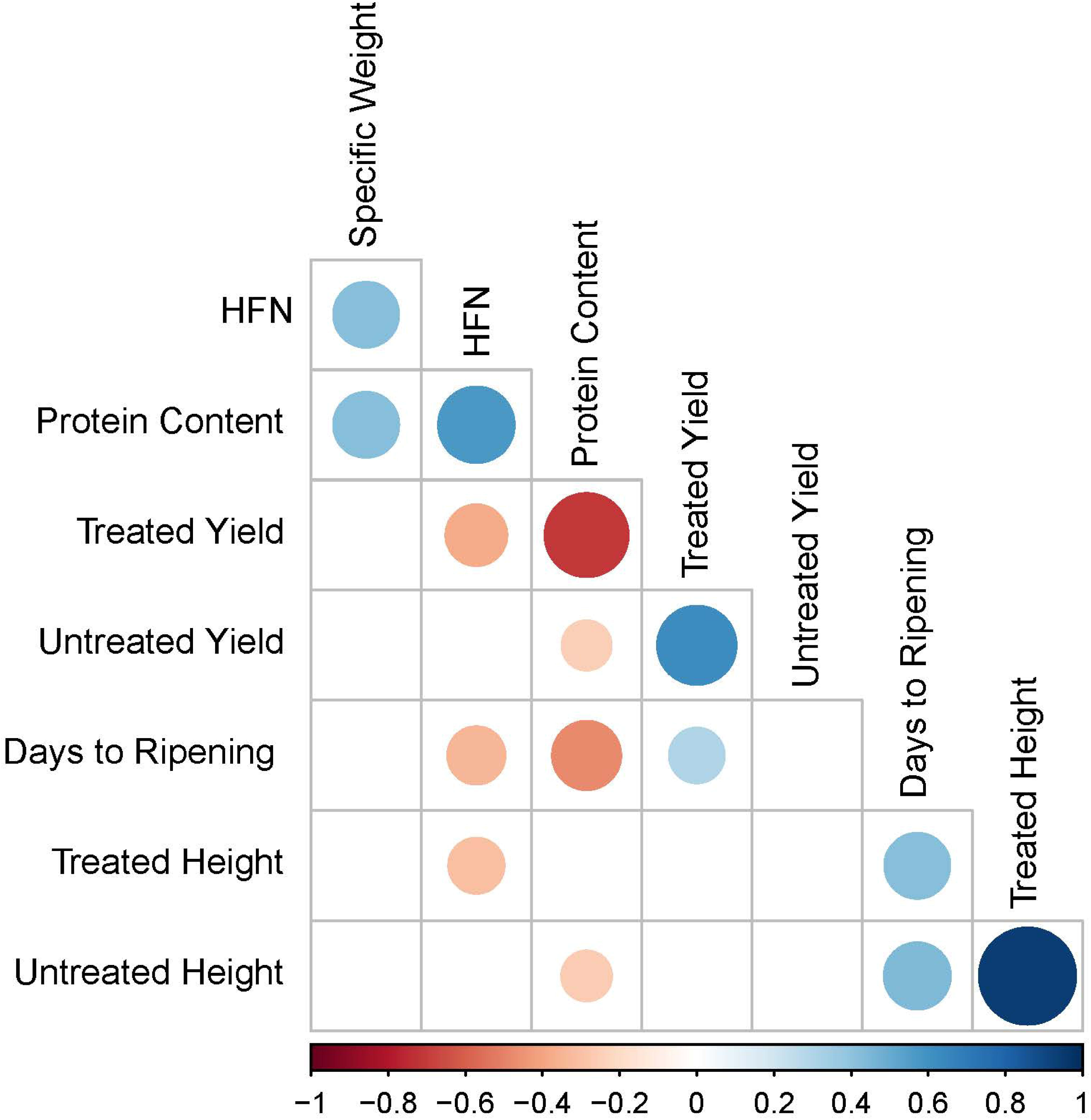
Phenotype correlation between yield, adaptation and grain quality traits. EVM were derived for each variety from the NVPT conducted between 2002 and 2017. Only significant correlations (P < 0.05) are indicated. Positive and negative correlations are indicated with the blue and red circles, respectively, with the size and colour intensity of the circles representing the magnitude of the correlation.

### Examining Trait Trends

We next examined the temporal pattern across the 15 years of trials to highlight linear trends in traits due to breeding progress. For this, we regressed the EVM for each variety on its year of first entry to the NVPT which is directly related to its year of release. This regression likely captures temporal pattern of breeding progress as successive releases of varieties are expected to outperform previous releases in one or more traits. We observed linear increase for yield between 2002 - 2017 in both the treated and untreated trials (Figure 3A - B). The rate of yield increase in the untreated trial was significantly higher than in the treated trials (rate difference = 0.093 tonnes/ha/year, P < 0.0001). Conversely, grain protein content and HFN showed small but significantly decrease over time (P < 0.001 and 0.03, respectively; Figure 3C - D). We also observed a significant delay in days to ripening over the same period (P = 0.004, Figure 3E). Changes in plant height (treated and untreated) and specific weight were not significant (P = 0.31 – 0.51, Figure 3F - H) suggesting stable trends.

**Figure 3:**
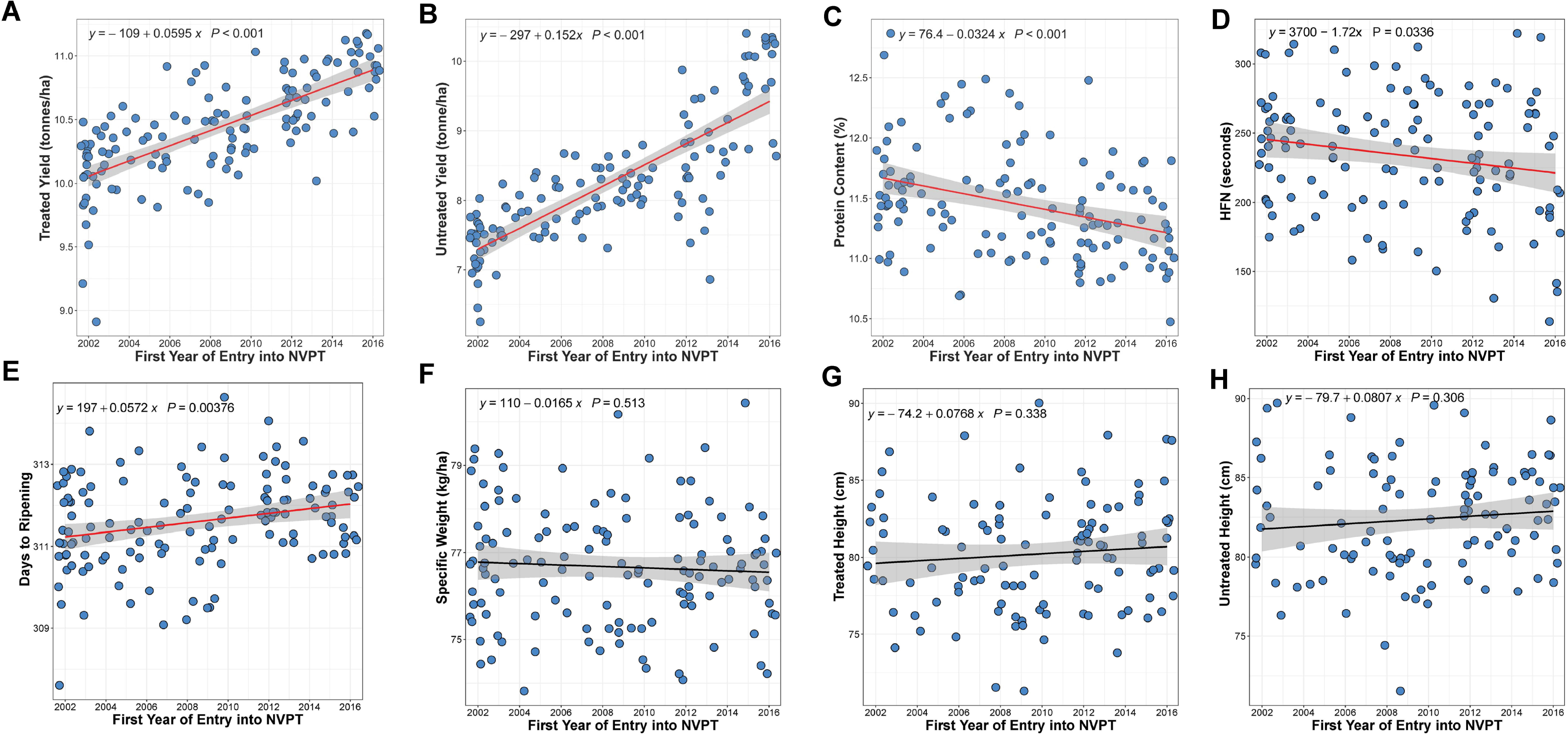
Temporal Trait Trend in UK Winter Wheat. Scatter plot showing changes in yield in the treated (A) and untreated trial (B), protein content (C), HFN (D), days to ripening (E), specific weight (F), plant height in treated (G) and untreated (H) trials. Blue dots represent individual varieties. For each trait, the EVM for each variety is regressed against the first year of entry in the 2002 −2017 trials. The solid line shows the regression line of the linear model and is coloured red if significant (P < 0.05). The shaded region defines the confidence interval. The regression equation is shown within each plot. The EVM data used for these plots are in Table S2.

UK wheat varieties are classified into four main end-use groups as described by the UK Flour Millers (www.ukflourmillers.org). These include the UK Flour Group 1 – 4, hereafter referred to as UFG1-4. The UFG1 and UFG2 varieties have superior grain quality (grain protein content and HFN) and are used for breadmaking. UFG3 varieties are often used for biscuits and cakes, whereas UFG4 varieties usually have high yield potential but inferior grain quality and are mainly used for animal feed. As yield and protein content are important measures for these end-use classifications, we examined how the temporal trends observed for these traits varied for the different end-use groups. Expectedly, UFG4 varieties showed higher yield while the bread making varieties (UFG1-2) show higher grain protein content (Figure 4A - B). All end-use groups showed a significant increasing yield trend across time and the rates of increase were not significantly different between the end-use groups (P = 0.263 − 0.885; Figure 4C). UFG2 and UFG4 varieties showed a significant and comparable decline in grain protein content over time (Figure 4D) while changes in protein content of UFG1 and UFG3 varieties were non-significant (Figure 4D).

**Figure 4:**
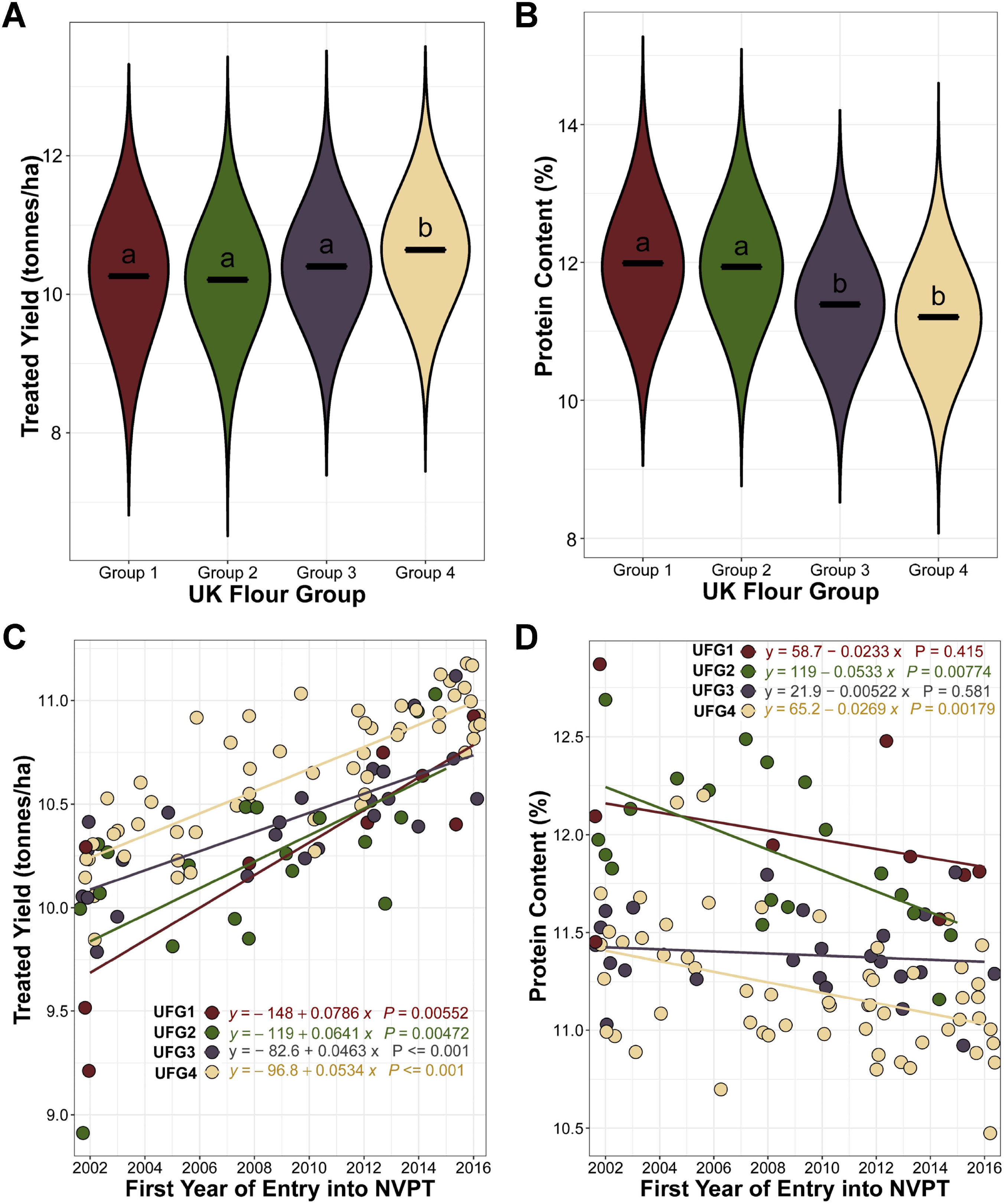
Temporal Trait Trend by End-use Groups. (A-B) Violin plots showing distribution for yield (A) and protein content (B) for the different end-use groups. The solid lines represent the mean of the distribution and the black letters show Tukey statistical comparison between the groups. Groups that are statistically similar share the same letter. (C-D) Scatter plot showing changes in yield (C) and protein content (D) for each end-use group of UK winter wheat. Each dot represents a variety while the colors 31 of the dots represent the end use groups (UK Flour Group 1-4). For each trait, the EVM for each variety is regressed against the first year of entry in the 2002 −2017 trials. The solid lines are the regression line of the linear model. The regression line equation for each group is shown. UFG1, UFG2, UFG3 and UFG4 are represented by the red, green, gray and peach dots, lines and text, respectively.

### Yield, protein content, specific weight, but not HFN, are stable in UK environments

Using a modified Finlay Wilkinson (FW) regression (Lian and de los Campos 2016) for measuring genotype x environment interaction, we examined the stability of yield and end-use quality traits across the trial years (Figure 5, Table S3). Only 95 varieties that were trialed in three or more years were included in this analysis. FW regression measures the stability of variety performance across different environments by regressing individual variety trait means on the environmental effect (Finlay and Wilkinson 1963). FW regression coefficient close to 1 suggests average varietal stability in which variety performance is consistent with environment effect i.e. variety performs poorly in bad environments and well in good environments. Larger values suggest below average stability i.e. higher environmental sensitivity.

**Figure 5:**
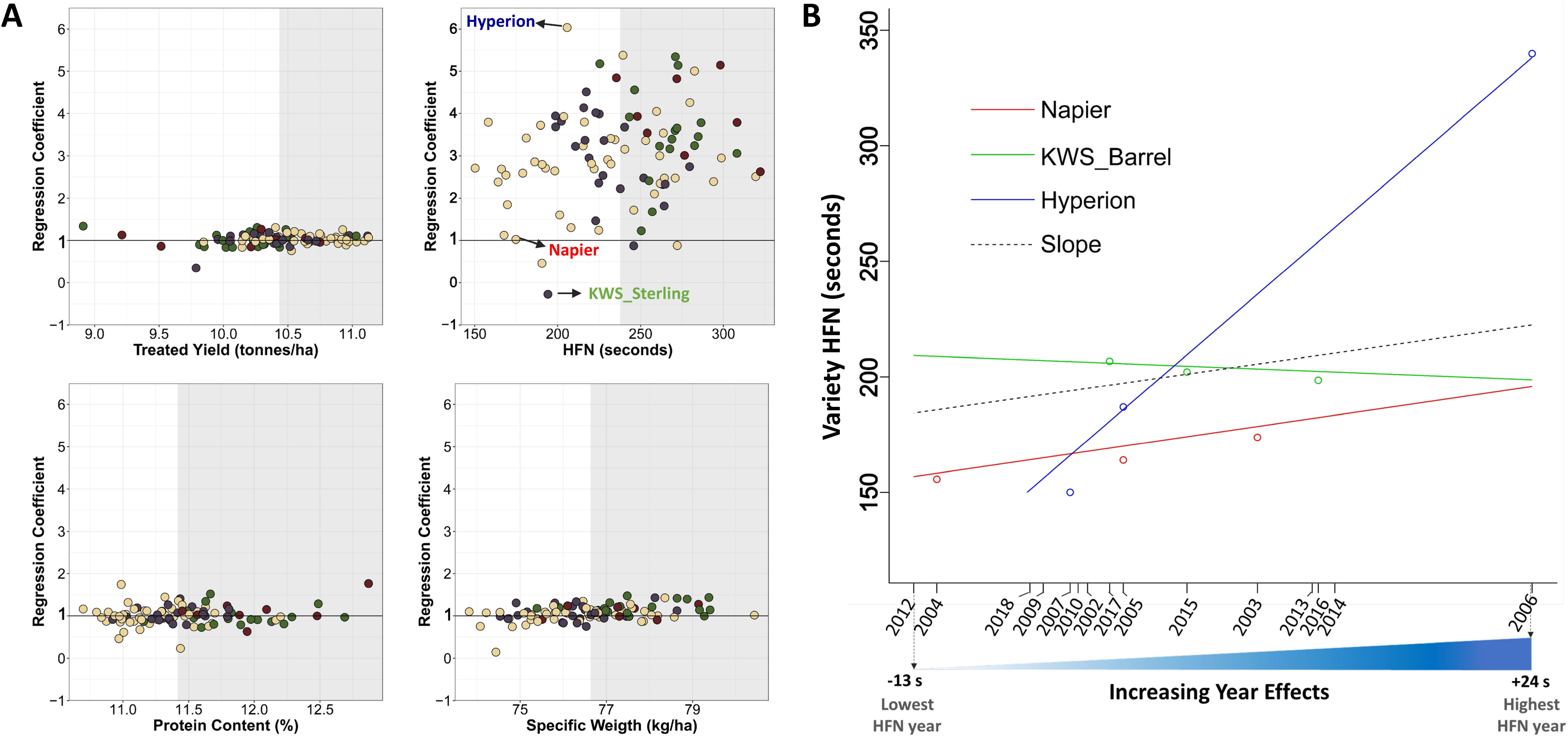
Phenotype Stability by End-use Group. Scatter plot showing stability for treated yield, protein content, specific weight and HFN for UK winter wheat varieties across the 15 years of trials (2002 – 2017). The y-axis represents the Finlay Wilkinson (FW) coefficient which specifies expected change in performance per unit change in environment (year) effect. Varieties with above median performance are in the shaded region. The solid line indicates stable performance in all environments i.e. b + 1 = 1 (Lian and De los Campos, 2016). Datapoints for three varieties whose HFN performance are further illustrated in Figure 5B are labeled. (B) Plot of HFN performance of varieties with lowest, highest and stable (~1) FW coefficient against the estimated environment year effect. The dashed lines present a constant slope of 1.

Yield was stable across years in most UK wheat varieties (regression coefficients close to 1, Figure 5A). Similarly, most of the varieties examined showed high stability in protein content and specific weight, with bread-making varieties stably producing grains with above median protein and specific weights (Figure 5B). HFN, on the other hand, showed varying FW coefficients ranging from −0.28 (KWS Barrel) to 6.03 (Hyperion). More than 83% of the 95 UK wheat varieties examined have FW coefficient > 2 for HFN suggesting below-average stability. Figure 5B shows the HFN performance of three varieties with different FW coefficients: KWS_Barrel, Hyperion, and Napier with (FW coefficient of 1.02). Napier consistently showed low HFN values in all the years it was trialed. On the other hand, Hyperion with the highest FW co-efficient, showed extreme HFN phenotypes - very low HFN value in Low-HFN years and very high HFN value in high-HFN years suggesting high environmental sensitivity. KWS_Barrel’s HFN performance was fairly constant irrespective of the environments it was trialed.

### Post-2002 UK wheat varieties belong into four distinct population groups

Using the Axiom35k SNP array (Allen *et al.* 2017) we genotyped 139 varieties including a subset of those trialed between 2002 - 2017 (104) and additional historic UK wheat cultivars. After quality filtering (described in Materials and Methods), we selected 4298 high quality markers dataset (Table S4) including 1715, 1781 and 778 markers on the A, B and D sub-genomes, respectively (Table S5). Using these genotypic data, we examined the population structure within the UK wheat collection. DAPC analysis revealed four distinct population groups (Pop1-4; Figure 6A, Table S6). Using Helium for pedigree visualization (Shaw *et al.* 2014), we could trace the modern founder parents for three (Pop1, 2 and 4) of the four population groups. Pop1 contains 19 varieties, of which 15 (79%) have Cadenza in their pedigree, consistent with Cadenza being an important parent for Pop1. Pop2 comprises 27 varieties, 20 (74%) of which contain Claire in their pedigree. Pop4 includes 30 varieties, 28 (93%) of which trace their pedigree to Robigus suggesting Robigus as an important parent for this group (Figure 6B). Pop3 is the largest group with 63 varieties with a more diverse pedigree structure. Using a subset of 111 varieties with both genotype and end-use group information (Figure S2), we examined the association between the population groups and end-use groups (Figure S3). The “Claire” (Pop2) and “Robigus” (Pop4) population groups only contain UFG3 and UFG4 varieties used for biscuit/cakes and feeds, respectively. While the “Cadenza” (Pop1) population group mostly (71%) contain UFG1 and UFG2 varieties used for breadmaking.

**Figure 6:**
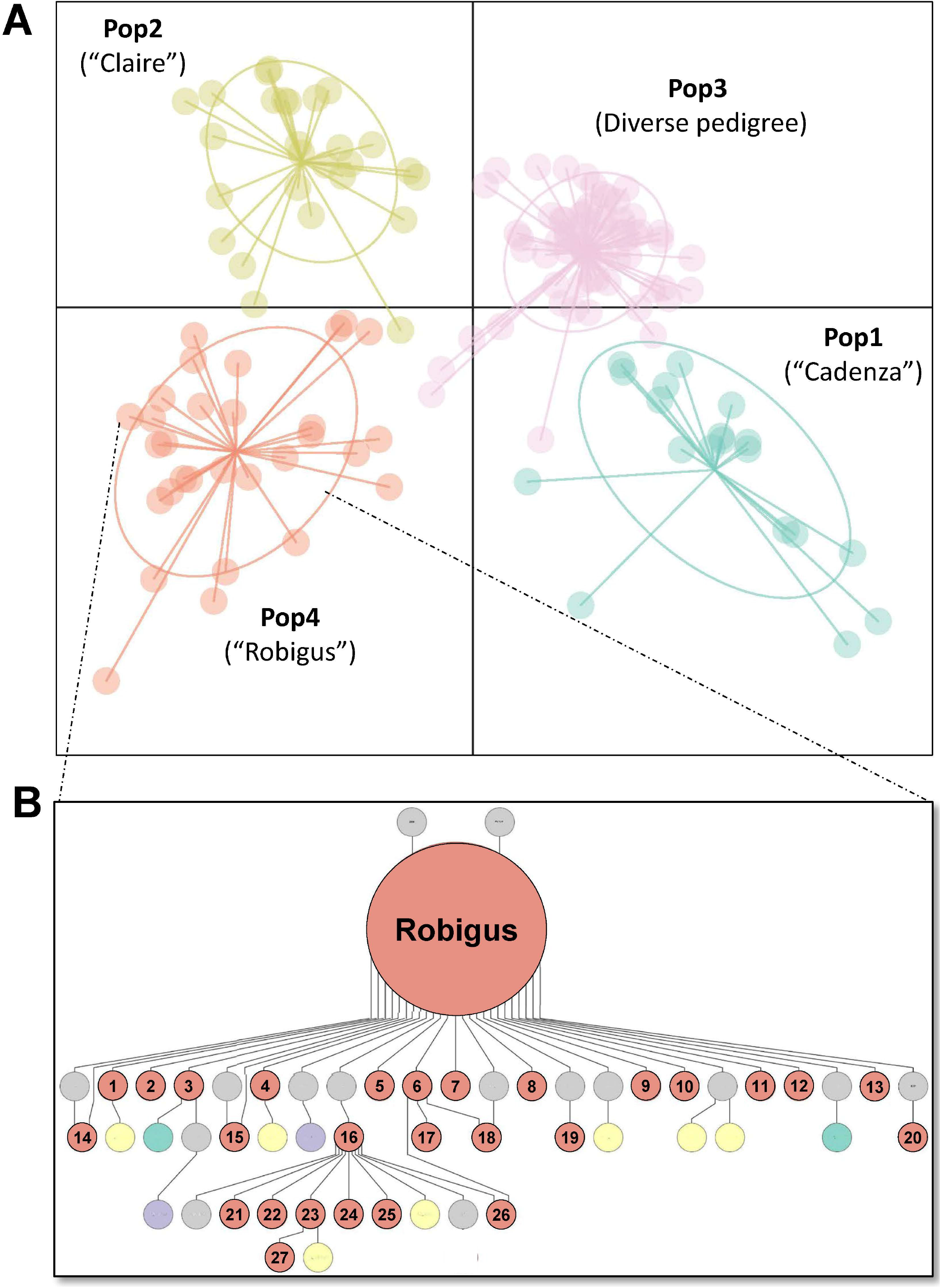
Population structure of UK winter wheat varieties using DAPC analysis. (A) The representative variety for each population group (Pop) is indicated except for Pop2 which consists of a more diverse pedigree. (B) Pedigree structure for Pop4 “Robigus”. The number in the inset represent varieties: (1) Qplus (2) Torch (3) Viscount (4) Conqueror (5) Leeds (6) Lear (7) Zulu (8) Gravitas (9) Twister (10) Britannia (11) Invicta (12) Warrior (13) Cougar (14) KWS Croft (15) KWS Target (16) Oakley (17) Jorvik (18) Panacea (19) Tuxedo (20) Icon (21) Horatio (22) KWS Gator (23) KWS Santiago (24) RGT Scrummage (25) Reflection (26) Energise (27) KWS Kerrin. The population groups are represented by teal (Pop1), yellow (Pop2), purple (Pop3), and red (Pop4) circles, whereas gray circles represent varieties which were not genotyped in this study.

### Using NVPT Data for Trait Mapping

We next examined the suitability of using the EVM obtained from the NVPT for trait mapping through a genome-wide association study (GWAS). To ascertain that our genotypic data and population composition are suitable for GWAS, we included data for the presence/absence of *Sm1* - a major locus known to underlie resistance to Orange wheat blossom midge (OWBM) in UK wheat varieties. As expected, we identified a major peak associated with OWBM resistance on wheat chromosome 2B (Figure S4A and B). This peak co-localizes with the physical position for *Sm1* (Walkowiak *et al.* 2020), supporting our *Sm1* marker information. Importantly, our GWAS analysis identified a region on the short arm of chromosome 6A with significant marker trait association (MTA) for days to ripening (Figure 7A - B). The days to ripening MTA region contain two markers, AX-94549511 and AX-94710688, located in an interval (73.5 – 86.5 Mbp) containing the *NAM-A1* gene (TraesCS6A02G108300; 77.1 Mbp) that is associated with variation in senescence in European wheat cultivars (Cormier *et al.* 2015). Days to ripening was significantly different between the allele groups of marker AX-94710688 which has the highest significance score (Figure 7C).

**Figure 7:**
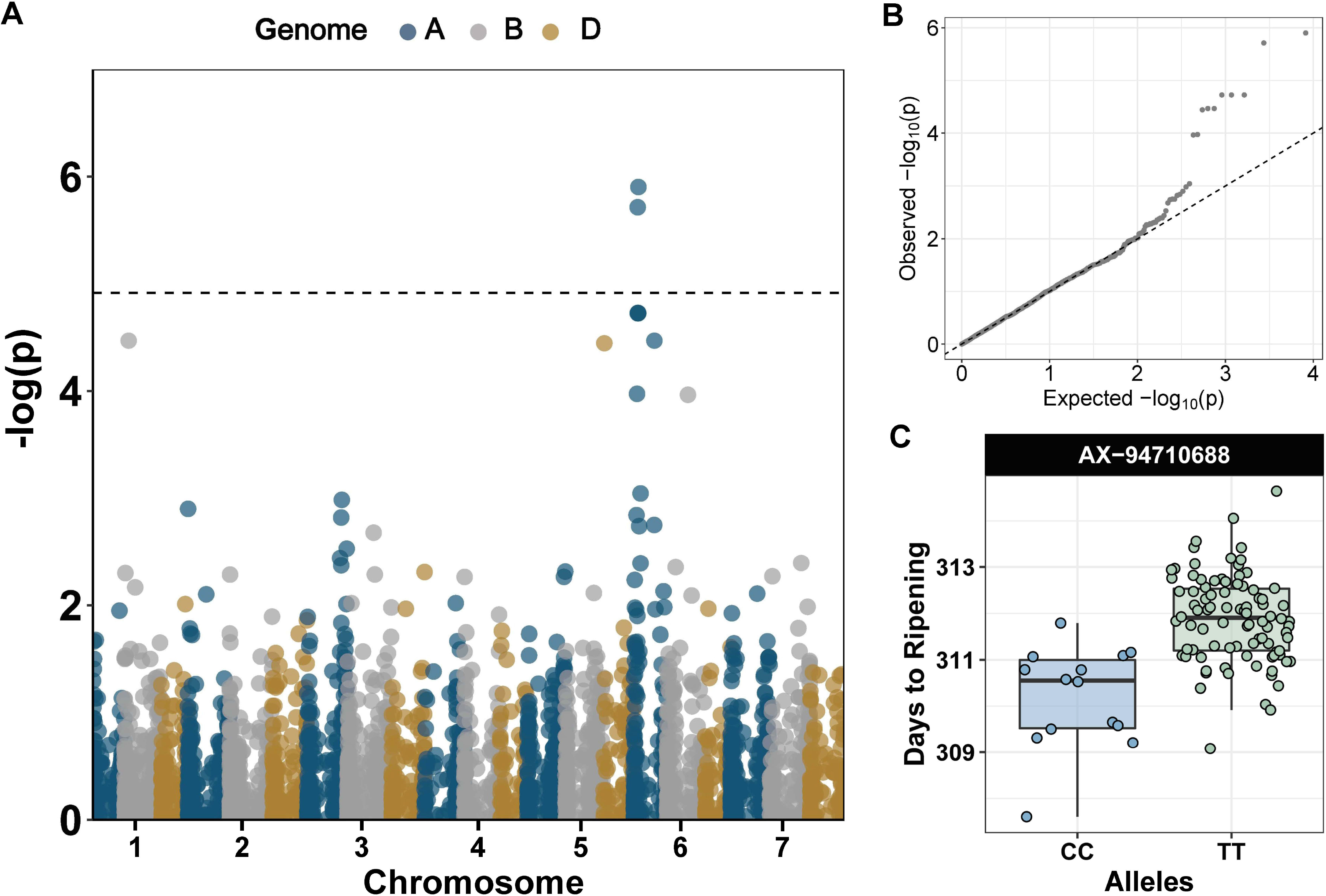
(A) Manhattan plot for days to ripening using EVM derived from the 2002 – 2017 NVPT of UK winter wheat varieties. The Bonferroni threshold is indicated with a dotted line. The seven wheat chromosome groups are indicated on the X-axis and each homoeologous sub-genome is coloured in red (A genome), gray (B) or yellow (D). (B) QQplot showing expected and observed distribution of –log (p values). (C) Allele effect of the marker showing the highest significant marker trait association for days to ripening.

## DISCUSSIONS

### Yield is an important driver of linear trends

Using historical data from UK NVPT we examined phenotypic trends in winter wheat varieties trialed between 2002 – 2017. Our analysis highlights a linear increase for yield (treated and untreated) and days to ripening, and a linear decrease in protein content and HFN. Given that the model used to analyze this data adjusted for variation arising from locations across years, and that agronomic practices are largely consistent in the NVPT, this linear trend can be attributed mostly to genetic improvement of varieties over time. Mackay *et al.* (2011) similarly attributed 88% of yield increase in cereals crops in the UK from 1982 – 2007 to genetic improvement. Yield is the most important determinant of grain market value; as such the linear increase in yield is consistent with concerted breeding efforts to improve yield under UK wheat growing conditions. In addition to the overall yield trend, we also observed consistent and similar linear increases in yield in all the four UK Flour Groups (UFG1 – 4). This further highlight yield as the main breeding target for varietal development (and adoption into the RL) irrespective of their target end-use groups.

We observed that the rate of yield increase in untreated trials (152 kg/ha/year) is significantly (p <0.0001) higher than in treated trials (60 kg/ha/year) across the 15-year period. Mackay *et al.* (2011) similarly observed the same pattern between 1982 – 2007 and argued that this pattern is due to loss of disease resistance by some varieties during the trial period examined. Varieties progressively lose resistance over time (Meikle and Scarisbrick 1994) and consequently variety performance declines with time. This mean that under untreated trial conditions, newly introduced varieties with ‘intact’ disease resistance will outperform a portion of previously released varieties whose disease resistance have ‘broken down’. This differential loss of disease resistance will further increase the variation in variety yield performance in untreated trials in addition to the variation arising from non-disease related genetic factors observed in treated trials. In other words, there is an “upward bias” in variety effects for the yield observed in untreated trials as described by Mackay *et al.* (2011).

Based on the rationale described above, it would be expected that a sudden loss of resistance in a large proportion of varieties due to the emergence of a more virulent pathogen race would result in a marked upward bias in variety effect estimates. This is what we observed when we compared yield trends before and after the emergence of the yellow rust (*Puccinia striiformis*) “Warrior” race in 2011 (Hubbard *et al.* 2015). The rate of yield increase in untreated trials significantly (P < 0.001) increased three-fold from 123 kg/ha/year before the emergence of the “Warrior” race to 372 kg/ha/year after the emergence of the “Warrior” race (Figure 8). During the same time, the rate of yield increase was significantly (P = 0.2697) comparable in the treated trial before and after the emergence of the “Warrior” race (Figure 8). The use of historical data in this study allowed us to identify this trend and thus highlight the importance of such datasets for dissecting the effect of important events in a national cropping history such as change in disease epidemics.

**Figure 8:**
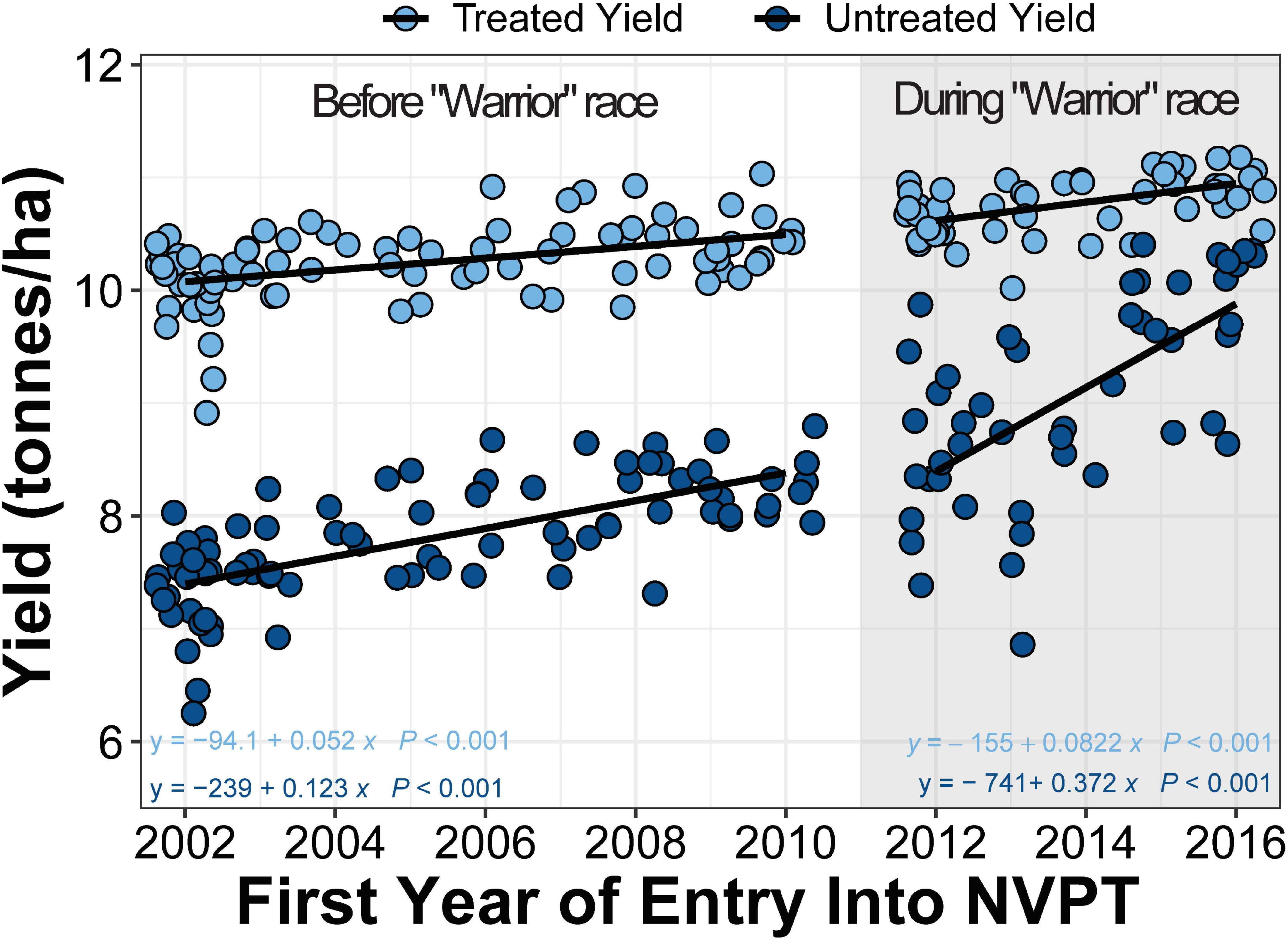
Yield comparison between treated and untreated trials before and after the emergence of the “warrior” yellow rust race. Scatter plot showing changes in yield in treated (light blue) and untreated trials (dark blue) before (unshaded region) and after (shaded region) the emergence of the “warrior” yellow rust race. The EVM for each period are regressed separately against the first year of entry into the NVPT trials for each variety. The solid lines are the regression lines. The regression equations are shown at the bottom corners of the plot.

It is also interesting to speculate that the higher rate of yield increase observed in the untreated trials indirectly suggests that newer varieties contain new sources of genetic resistance that improve their performance over older varieties at a rate greater than observed in the treated trials. This is likely not accidental, but points to concerted efforts by breeders to introduce more effective source of genetic resistance into UK wheat. The improved genetic resistance profile of newer varieties narrows the yield gap observed between the treated and untreated trials. We cannot, however, rule out the fact that this narrower yield gap might be due to less disease pressure in recent years. A more detailed genetic characterization will be needed to accurately describe the genetic resistance profile of UK wheat varieties.

Concomitant with the yield increase, there has been a decrease in grain protein content from 2002 - 2017 which reflects the well-established antagonistic relationship between yield and protein content (Figure S5; Simmonds 1995). Unlike for yield, linear trends were not consistent across the four end-use groups. While we identified an overall significant decrease in grain protein content over time, this was not observed in the UFG1 varieties that are used for breadmaking (Figure 4). UFG2 varieties which also have breadmaking potential, however, showed significant decrease over time just like the UFG4 varieties used for animal feed. The decline in UFG2 varieties grain protein content may be due to the fact that this group comprise varieties that did not consistently meet the higher grain quality (in particular protein content) requirement for UFG1 and were downgraded to UFG2. The fact that our analysis captures expected trait (yield, protein content and HFN) differences in end-use groups (Figure 4A - B, Figure S6A - B) suggests that the linear mixed effect model adopted is appropriate to handle the incomplete design of the NVPT and to examine phenotype trends within each end-use group.

The multidimensional (year and location) nature of the NVPT also allows for examining varietal adaptability across multiple environments. We observed year-to-year stability in yield and protein content in most of the varieties irrespective of their end-use group. This is likely attributable to the fact that we mainly examined data from RL trials that are comprised of varieties which had been previously screened for distinctness, uniformity, and stability during National Listing trials. Despite this ‘pre-screening’, almost all the varieties show high environmental sensitivity for HFN (FW coefficient: −0.28 to 6.03). Sjoberg et al (2020) similarly obtained a wide range of FW coefficient for HFN in 133 varieties trialed across three years in the Pacific Northwest of the US.

HFN is inversely related to α-amylase activity within the grain. High α-amylase activity caused by incidences of pre-harvest sprouting (PHS) and/or pre-maturity amylase (PMA) reduce the bread-making potential of wheat grains. Both PHS and PMA are known to be highly environmental dependent: PHS is induced by wet raining conditions during harvest maturity while PMA is mostly caused by low or high temperature shock around grain physiological maturity (Joe *et al.* 2005; Mares and Mrva 2014). The environmental conditions required to induce PHS and PMA occur infrequently from year to year making it difficult for breeders to screen for these traits under field conditions. In addition, both traits are controlled by many genes most of which have small effects making marker assisted selection (MAS) for HFN stability difficult. Within the last decade, progress has been made in identifying genes with major effects on PHS including *TaMFT* and *TaMKK3*-A (Nakamura *et al.* 2011; Torada *et al.* 2016). We also previously showed the effect of *TaMMK3*-*A* in reducing PHS in UK germplasm (Shorinola *et al.* 2016) and developed markers to facilitates its use in breeding (Shorinola *et al.* 2017). The availability of markers for major genes controlling PHS now makes it possible to apply MAS for improving HFN. However, selection for PMA resistance remains a major challenge because the conditions that induces PMA varies between varieties (Liu *et al.* 2021)

### Population structure within UK winter wheat germplasm

Our analysis reveals that three modern wheat varieties largely contribute to the development of winter wheat varieties released in the UK between 2002 - 2017. These include Cadenza (Pop1), Claire (Pop2) and Robigus (Pop4), which were themselves released in 1992, 1999, 2005, respectively. Together, 51% of the 114 varieties that were first trialed between 2002 – 2017 were derived from either Cadenza, Claire, and/or Robigus. Based on pedigree visualization, Robigus (and Pop4 varieties) appears to be a more recent introduction to the UK (Figure S7) suggesting that new gene pools are being introduced into the UK wheat breeding landscape. Since its introduction Robigus has made significant contribution to UK wheat pedigree. Fradgley et al (2019) identified Robigus as the second most used parents in UK breeding, next to Capelle Desprez. We also observed a clear association between the population groups and end use groups. Pop2 and Pop4 varieties, mostly derived from Claire and Robigus which are themselves UFG3 varieties, both contain only UFG3 (biscuit) and UFG4 (feed) varieties. Pop1 varieties, which are mostly derived from Cadenza - a UFG2 variety, mostly contain UFG1 and UFG2 (breadmaking) varieties. One probable explanation for this association is that breeders tend to make crosses with varieties from the same end-use groups to ensure that the gene combinations underlying the traits in the target end-use groups are preserved in their progenies (Simon Berry 2021, personal communication). This suggests that the choice of parents is an important determinant of the end-use class of varieties.

Due to the type (gene-based SNP) and limited number of markers used, we acknowledge the limitation of this study to more precisely define the population groups represented in UK winter bread wheat collection to a high resolution. Brinton et al. (2020) demonstrated the inadequacy of array-based genotyping chips to precisely define haplotype groups due to their gene-centric design. Scaffold-level assemblies are now available for important UK wheat varieties including representatives of Pop1, Pop3 and Pop4 (Cadenza, Claire and Robigus; Walkowiak *et al.* 2020). These genome assemblies can be combined with high-density genotyping or re-sequencing data to more precisely define the populations groups of wheat varieties grown in the UK.

### Historical data could be valuable for trait mapping

We identified significant marker-trait association (MTA) peaks spanning a gene (*NAM-A1*) that have been previously associated with natural variation in a trait of agronomic interest.. Cormier et al. (2015) identified a C/T missense SNP in the NAC domain and A/- frame-shift deletion in *NAM-A1* leading to a truncated protein from a worldwide wheat collection and suggested functional roles for these polymorphisms. Harrington et al., (2019) showed that missense mutations in the NAC domain of *NAM-A1* result in delayed peduncle and flag leaf senescence. Similarly, Avni et al (2014) showed that loss of function *NAM-A1* mutants showed significant delay in senescence. Given the large interval covered by the MTA peaks for days to ripening on chromosome 6A (73.4 Mbp – 86.5 Mbp, ~140 genes) we cannot rule out the possibility that other gene(s) underly this days to ripening effect. Nonetheless, the co-localization of our GWAS peak with a known locus for the target trait highlights the usefulness of this historical dataset for quantitative trait mapping.

Beside the MTA for days to ripening, we did not identify strong MTA for the other traits. This might be due to the fact that many of the major genes controlling these traits have been mostly fixed in the UK wheat population examined, and that the population size used in our study is not large enough to pick up minor effect and/or minor allele frequency gene(s). Also, while the phenotyping conditions used in the NPVT might be representative of UK farming conditions, they might not always be best suited for trait mapping. An example is the application of plant growth regulators in the trials to prevent lodging (by reducing plant height) but this might mask the effect of height genes. Despite these limitations, our work demonstrates that national trials data can be valuable for examining trait trends, stability, and genetic architecture.

## Acknowledgments

We thank the Agriculture and Horticulture Development Board (AHDB) for making the trial data publicly available. We also thank Dr Chinyere Ekine for useful discussion on statistical analyses and Dr Simon Berry for suggestions to the manuscript.

## REFERENCES

Adamski, N. M., P. Borrill, J. Brinton, S. A. Harrington, C. Marchal et al., 2020 A roadmap for gene functional characterisation in crops with large genomes: Lessons from polyploid wheat (C. S. Hardtke, Ed.). eLife 9: e55646.

Allen, A. M., M. O. Winfield, A. J. Burridge, R. C. Downie, H. R. Benbow et al., 2017 Characterization of a Wheat Breeders’ Array suitable for high-throughput SNP genotyping of global accessions of hexaploid bread wheat (Triticum aestivum). Plant Biotechnology Journal 15: 390–401.

Avni, R., R. Zhao, S. Pearce, Y. Jun, C. Uauy et al., 2014 Functional characterization of GPC-1 genes in hexaploid wheat. Planta 239: 313–324.

Ayalew, H., P. W. Tsang, C. Chu, J. Wang, S. Liu et al., 2019 Comparison of TaqMan, KASP and rhAmp SNP genotyping platforms in hexaploid wheat. PLOS ONE 14: e0217222.

Bolser, D., D. M. Staines, E. Pritchard, and P. Kersey, 2016 Ensembl Plants: Integrating Tools for Visualizing, Mining, and Analyzing Plant Genomics Data, pp. 115–140 in Plant Bioinformatics: Methods and Protocols, edited by D. Edwards. Methods in Molecular Biology, Springer, New York, NY.

Brinton, J., R. H. Ramirez-Gonzalez, J. Simmonds, L. Wingen, S. Orford et al., 2020 A haplotype-led approach to increase the precision of wheat breeding. Communications Biology 3: 712.

Cheng, H., J. Liu, J. Wen, X. Nie, L. Xu et al., 2019 Frequent intra- and inter-species introgression shapes the landscape of genetic variation in bread wheat. Genome Biology 20: 136.

Cormier, F., M. Throude, C. Ravel, J. L. Gouis, M. Leveugle et al., 2015 Detection of NAM-A1 Natural Variants in Bread Wheat Reveals Differences in Haplotype Distribution between a Worldwide Core Collection and European Elite Germplasm. Agronomy 5: 143–151.

Crossa, J., J. Burgueño, S. Dreisigacker, M. Vargas, S. A. Herrera-Foessel et al., 2007 Association Analysis of Historical Bread Wheat Germplasm Using Additive Genetic Covariance of Relatives and Population Structure. Genetics 177: 1889–1913.

Finlay, K., and G. Wilkinson, 1963 The analysis of adaptation in a plant-breeding programme. Aust. J. Agric. Res. 14: 742–754.

Fradgley, N., K. A. Gardner, J. Cockram, J. Elderfield, J. M. Hickey et al., 2019 A large- scale pedigree resource of wheat reveals evidence for adaptation and selection by breeders. PLOS Biology 17: e3000071.

Gardiner, L.-J., L. U. Wingen, P. Bailey, R. Joynson, T. Brabbs et al., 2019 Analysis of the recombination landscape of hexaploid bread wheat reveals genes controlling recombination and gene conversion frequency. Genome Biology 20: 69.

Harrington, S. A., L. E. Overend, N. Cobo, P. Borrill, and C. Uauy, 2019 Conserved residues in the wheat (Triticum aestivum) NAM-A1 NAC domain are required for protein binding and when mutated lead to delayed peduncle and flag leaf senescence. BMC Plant Biology 19: 407.

He, F., R. Pasam, F. Shi, S. Kant, G. Keeble-Gagnere et al., 2019 Exome sequencing highlights the role of wild-relative introgression in shaping the adaptive landscape of the wheat genome. Nature Genetics 51: 896–904.

Howe, K. L., B. Contreras-Moreira, N. De Silva, G. Maslen, W. Akanni et al., 2020 Ensembl Genomes 2020—enabling non-vertebrate genomic research. Nucleic Acids Research 48: D689–D695.

Hubbard, A., C. M. Lewis, K. Yoshida, R. H. Ramirez-Gonzalez, C. de Vallavieille-Pope et al., 2015 Field pathogenomics reveals the emergence of a diverse wheat yellow rust population. Genome Biology 16: 23.

IWGSC, R. Appels, K. Eversole, N. Stein, C. Feuillet et al., 2018 Shifting the limits in wheat research and breeding using a fully annotated reference genome. Science 361: eaar7191.

Joe, A. F. T. W., R. W. Summers, G. D. Lunn, M. D. Atkinson, and P. S. Kettlewell, 2005 Pre-maturity α-amylase and incipient sprouting in UK winter wheat, with special reference to the variety Rialto. Euphytica 143: 265–269.

Jombart, T., and I. Ahmed, 2011 adegenet 1.3-1: new tools for the analysis of genome- wide SNP data. Bioinformatics 27: 3070–3071.

Jordan, K. W., S. Wang, Y. Lun, L.-J. Gardiner, R. MacLachlan et al., 2015 A haplotype map of allohexaploid wheat reveals distinct patterns of selection on homoeologous genomes. Genome Biology 16: 48.

Krasileva, K. V., H. A. Vasquez-Gross, T. Howell, P. Bailey, F. Paraiso et al., 2017 Uncovering hidden variation in polyploid wheat. Proc Natl Acad Sci USA 114: E913.

Lenth, R. V., 2016 Least-Squares Means: The R Package lsmeans. Journal of Statistical Software 69: 33.

Lian, L., and G. de los Campos, 2016 FW: An R Package for Finlay–Wilkinson Regression that Incorporates Genomic/Pedigree Information and Covariance Structures Between Environments. G3: Genes|Genomes|Genetics 6: 589.

Liu, C., K. M. Tuttle, K. A. Garland Campbell, M. O. Pumphrey, and C. M. Steber, 2021 Investigating conditions that induce late maturity alpha-amylase (LMA) using Northwestern US spring wheat (Triticum aestivum L.). Seed Science Research 1–9.

Mackay, I., A. Horwell, J. Garner, J. White, J. McKee et al., 2011 Reanalyses of the historical series of UK variety trials to quantify the contributions of genetic and environmental factors to trends and variability in yield over time. Theor Appl Genet 122: 225–238.

Mares, D. J., and K. Mrva, 2014 Wheat grain preharvest sprouting and late maturity alpha-amylase. Planta 240: 1167–1178.

Meikle, S. M., and D. H. Scarisbrick, 1994 Cereal Breeding and Varietal Testing. British Food Journal 96: 11–16.

Nakamura, S., F. Abe, H. Kawahigashi, K. Nakazono, A. Tagiri et al., 2011 A Wheat Homolog of MOTHER OF FT AND TFL1 Acts in the Regulation of Germination. Plant Cell 23: 3215–3229.

Pozniak, C. J., J. M. Clarke, and F. R. Clarke, 2012 Potential for detection of marker– trait associations in durum wheat using unbalanced, historical phenotypic datasets. Mol Breeding 30: 1537–1550.

Rasheed, A., and X. Xia, 2019 From markers to genome-based breeding in wheat. Theoretical and Applied Genetics 132: 767–784.

Rosyara, U. R., W. S. De Jong, D. S. Douches, and J. B. Endelman, 2016 Software for Genome-Wide Association Studies in Autopolyploids and Its Application to Potato. Plant Genome 9:.

Scott, M. F., N. Fradgley, A. R. Bentley, T. Brabbs, F. Corke et al., 2020 Limited haplotype diversity underlies polygenic trait architecture across 70 years of wheat breeding. bioRxiv 2020.09.15.296533.

Semagn, K., R. Babu, S. Hearne, and M. Olsen, 2014 Single nucleotide polymorphism genotyping using Kompetitive Allele Specific PCR (KASP): overview of the technology and its application in crop improvement. Molecular Breeding 33: 1–14.

Semenov, M. A., 2009 Impacts of climate change on wheat in England and Wales. Journal of The Royal Society Interface 6: 343–350.

Shaw, P. D., M. Graham, J. Kennedy, I. Milne, and D. F. Marshall, 2014 Helium: visualization of large scale plant pedigrees. BMC Bioinformatics 15: 259.

Shorinola, O., B. Balcárková, J. Hyles, J. F. G. Tibbits, M. J. Hayden et al., 2017 Haplotype Analysis of the Pre-harvest Sprouting Resistance Locus Phs-A1 Reveals a Causal Role of TaMKK3-A in Global Germplasm. Front Plant Sci 8: 1555–1555.

Shorinola, O., N. Bird, J. Simmonds, S. Berry, T. Henriksson et al., 2016 The wheat Phs-A1 pre-harvest sprouting resistance locus delays the rate of seed dormancy loss and maps 0.3 cM distal to the PM19 genes in UK germplasm. Journal of Experimental Botany 67: 4169–4178.

Silvey, V., 1981 The contribution of new wheat, barley and oat varieties to increasing yield in England and Wales 1947–78.

Simmonds, N. W., 1995 The relation between yield and protein in cereal grain. Journal of the Science of Food and Agriculture 67: 309–315.

Sjoberg, S. M., A. H. Carter, C. M. Steber, and K. A. Garland‐Campbell, 2020 Unraveling complex traits in wheat: Approaches for analyzing genotype × environment interactions in a multienvironment study of falling numbers. Crop Science 60: 3013–3026.

Su, G., P. Madsen, M. S. Lund, D. Sorensen, I. R. Korsgaard et al., 2006 Bayesian analysis of the linear reaction norm model with unknown covariates. J Anim Sci 84: 1651–1657.

Sweeney, D. W., J. Sun, E. Taagen, and M. E. Sorrells, 2019 Genomic Selection in Wheat, pp. 273–302 in Applications of Genetic and Genomic Research in Cereals, edited by T. Miedaner and V. Korzun. Woodhead Publishing.

Torada, A., M. Koike, T. Ogawa, Y. Takenouchi, K. Tadamura et al., 2016 A Causal Gene for Seed Dormancy on Wheat Chromosome 4A Encodes a MAP Kinase Kinase. Curr Biol 26: 782–787.

Trnka, M., S. Feng, M. A. Semenov, J. E. Olesen, K. C. Kersebaum et al., 2019 Mitigation efforts will not fully alleviate the increase in water scarcity occurrence probability in wheat-producing areas. Sci Adv 5: eaau2406.

Walkowiak, S., L. Gao, C. Monat, G. Haberer, M. T. Kassa et al., 2020 Multiple wheat genomes reveal global variation in modern breeding. Nature 588: 277–283.

Wang, S., D. Wong, K. Forrest, A. Allen, S. Chao et al., 2014 Characterization of polyploid wheat genomic diversity using a high-density 90 000 single nucleotide polymorphism array. Plant Biotechnology Journal 12: 787–796.

Wilkinson, P. A., A. M. Allen, S. Tyrrell, L. U. Wingen, X. Bian et al., 2020 CerealsDB— new tools for the analysis of the wheat genome: update 2020. Database 2020:.

Winfield, M. O., A. M. Allen, A. J. Burridge, G. L. A. Barker, H. R. Benbow et al., 2016 High-density SNP genotyping array for hexaploid wheat and its secondary and tertiary gene pool. Plant Biotechnology Journal 14: 1195–1206.

Yang, W., H. Feng, X. Zhang, J. Zhang, J. H. Doonan et al., 2020 Crop Phenomics and High-Throughput Phenotyping: Past Decades, Current Challenges, and Future Perspectives. Molecular Plant 13: 187–214.

