## Supplemental Figures for "Trend, Population Structure and Trait Mapping from 15 Years of National Varietal Trials of UK Winter Wheat"

### Slide 1
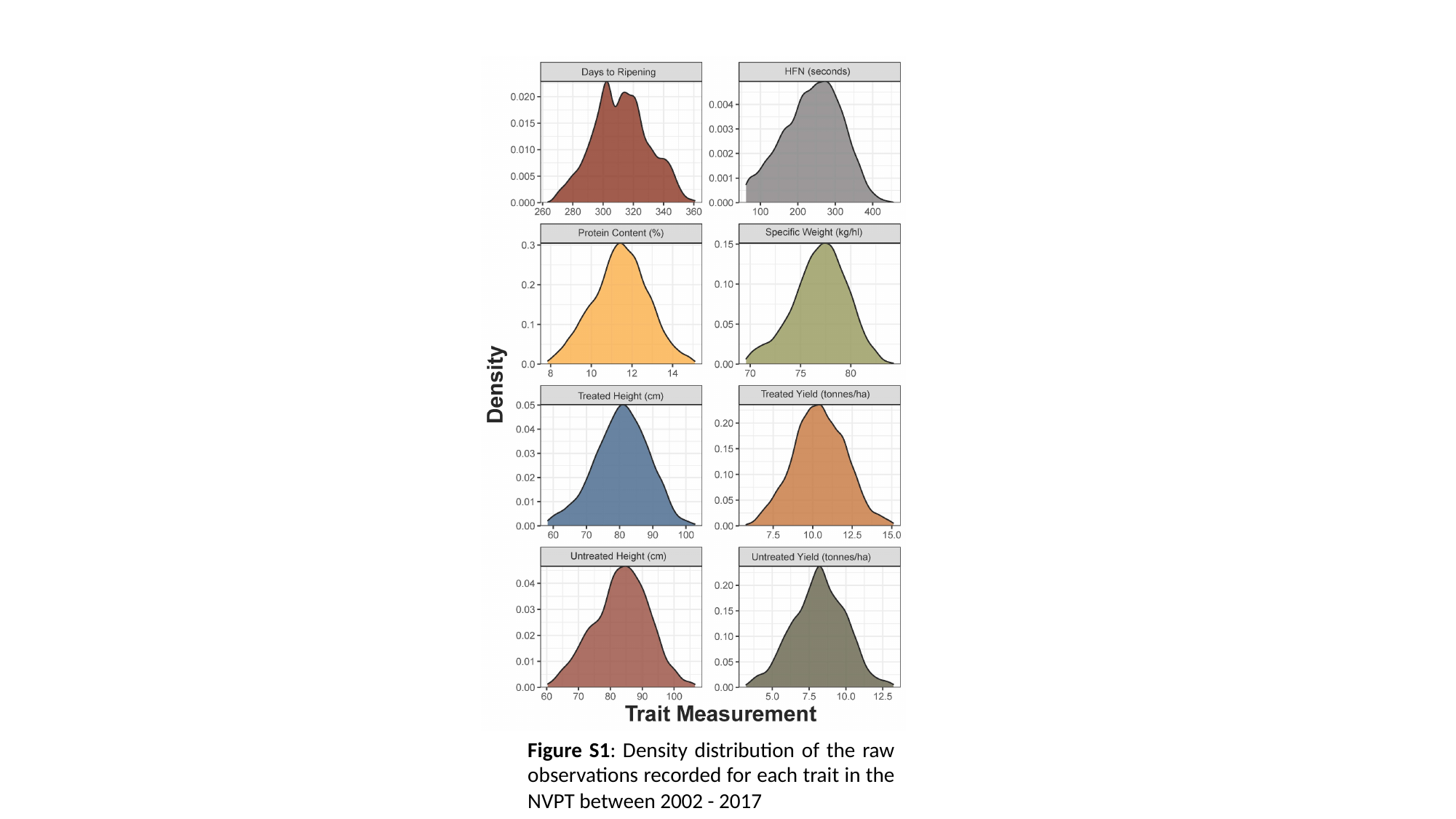

Figure S1: Density distribution of the raw observations recorded for each trait in the NVPT between 2002 - 2017

### Slide 2
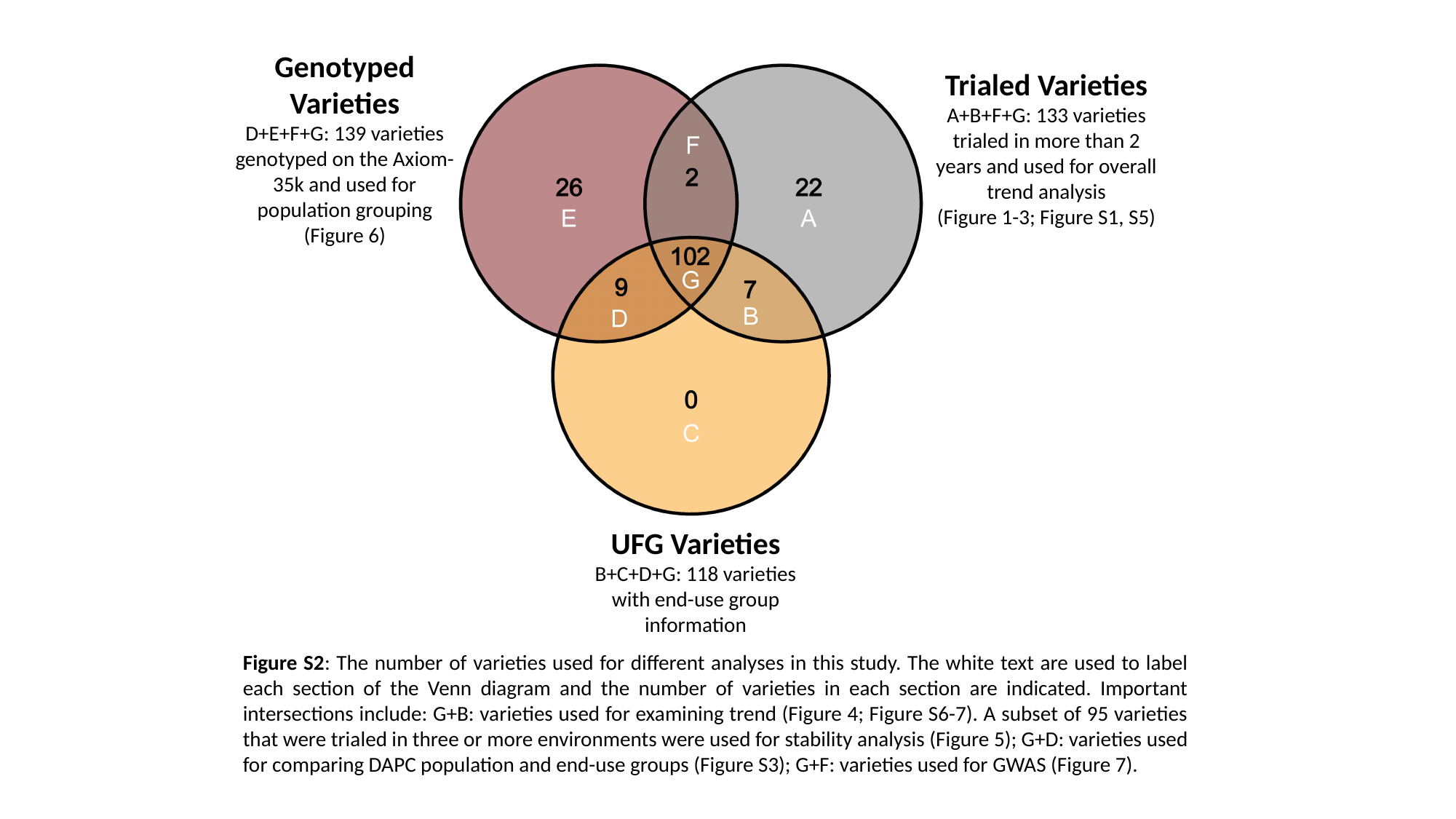

Genotyped Varieties
D+E+F+G: 139 varieties genotyped on the Axiom-35k and used for population grouping
(Figure 6)
Trialed Varieties
A+B+F+G: 133 varieties trialed in more than 2 years and used for overall trend analysis
(Figure 1-3; Figure S1, S5)
UFG Varieties
B+C+D+G: 118 varieties with end-use group information
Figure S2: The number of varieties used for different analyses in this study. The white text are used to label each section of the Venn diagram and the number of varieties in each section are indicated. Important intersections include: G+B: varieties used for examining trend (Figure 4; Figure S6-7). A subset of 95 varieties that were trialed in three or more environments were used for stability analysis (Figure 5); G+D: varieties used for comparing DAPC population and end-use groups (Figure S3); G+F: varieties used for GWAS (Figure 7).

### Slide 3
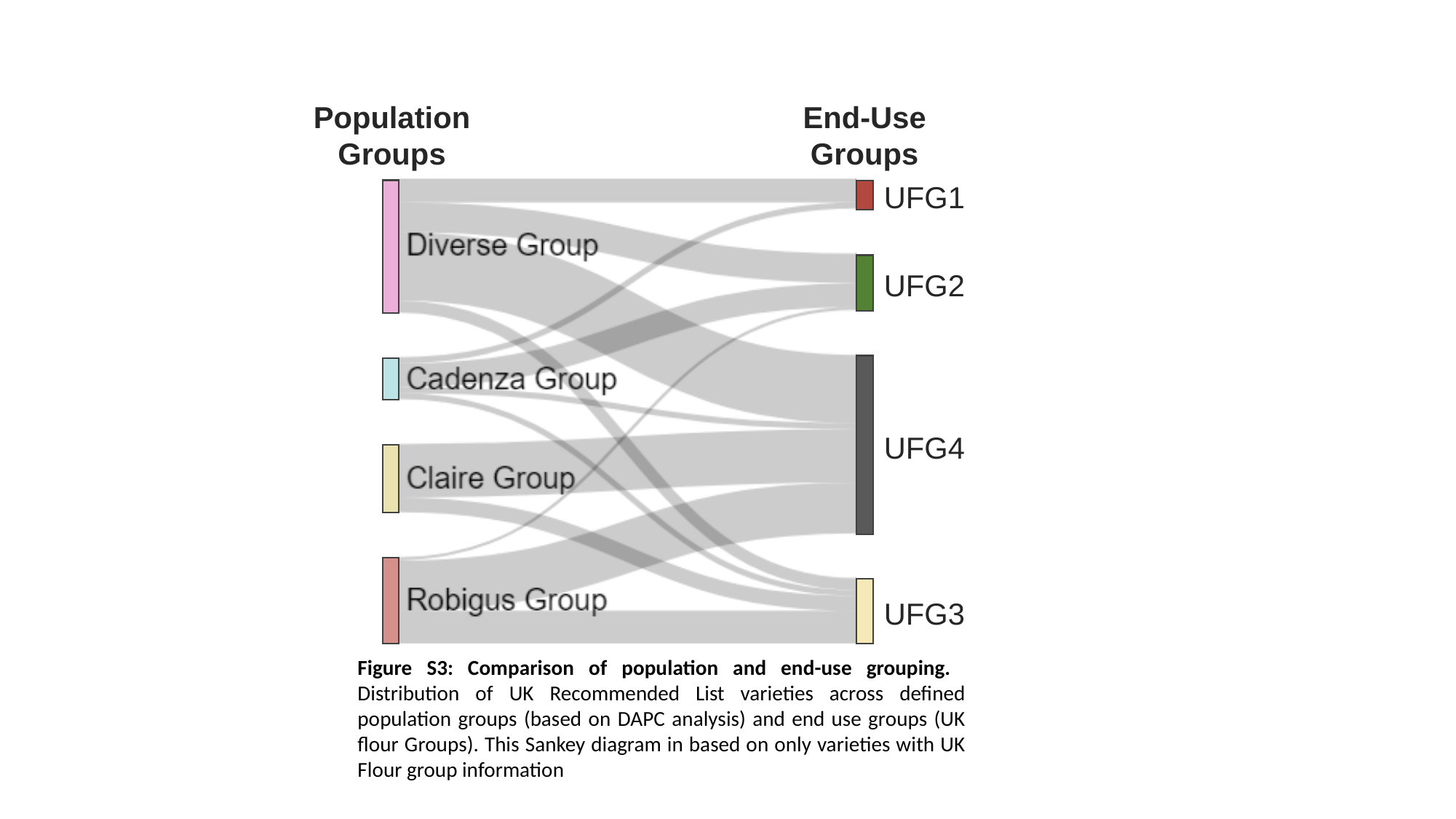

Population Groups
End-Use Groups
UFG1
UFG2
UFG4
UFG3
Figure S3: Comparison of population and end-use grouping. Distribution of UK Recommended List varieties across defined population groups (based on DAPC analysis) and end use groups (UK flour Groups). This Sankey diagram in based on only varieties with UK Flour group information

### Slide 4
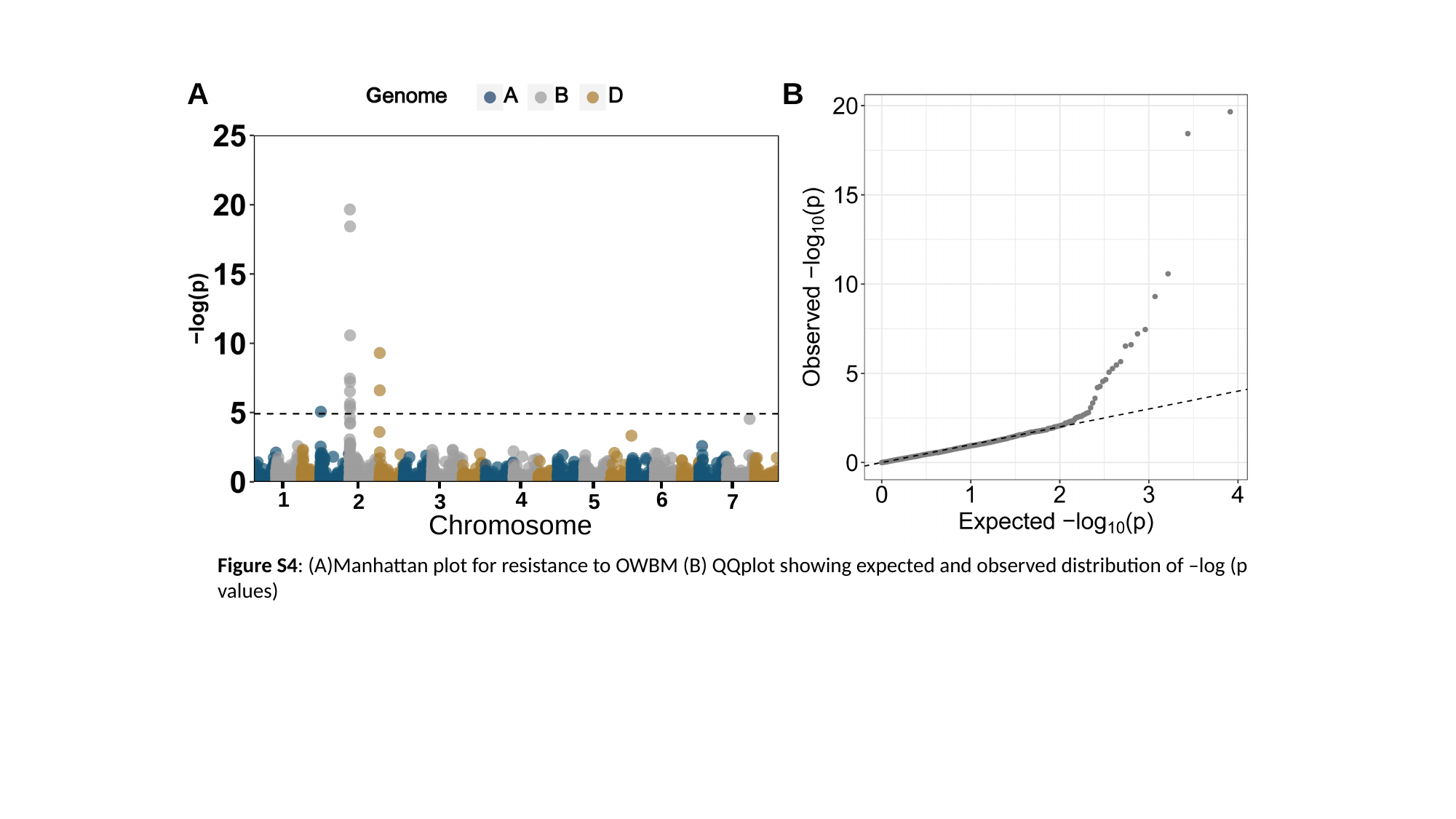

A
B
1
4
6
2
3
5
7
Chromosome
Figure S4: (A)Manhattan plot for resistance to OWBM (B) QQplot showing expected and observed distribution of –log (p values)

### Slide 5
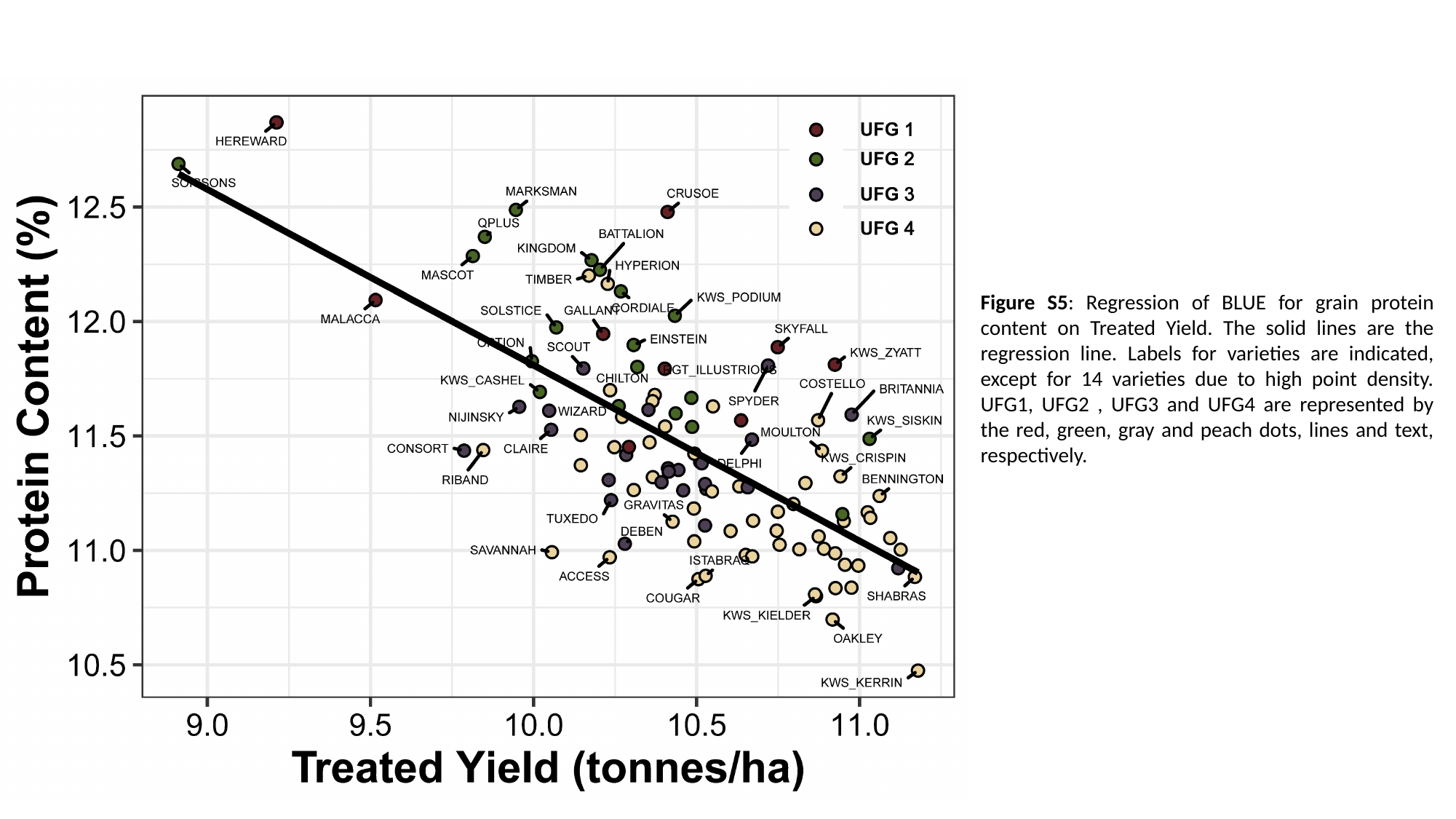

Figure S5: Regression of BLUE for grain protein content on Treated Yield. The solid lines are the regression line. Labels for varieties are indicated, except for 14 varieties due to high point density. UFG1, UFG2 , UFG3 and UFG4 are represented by the red, green, gray and peach dots, lines and text, respectively.

### Slide 6
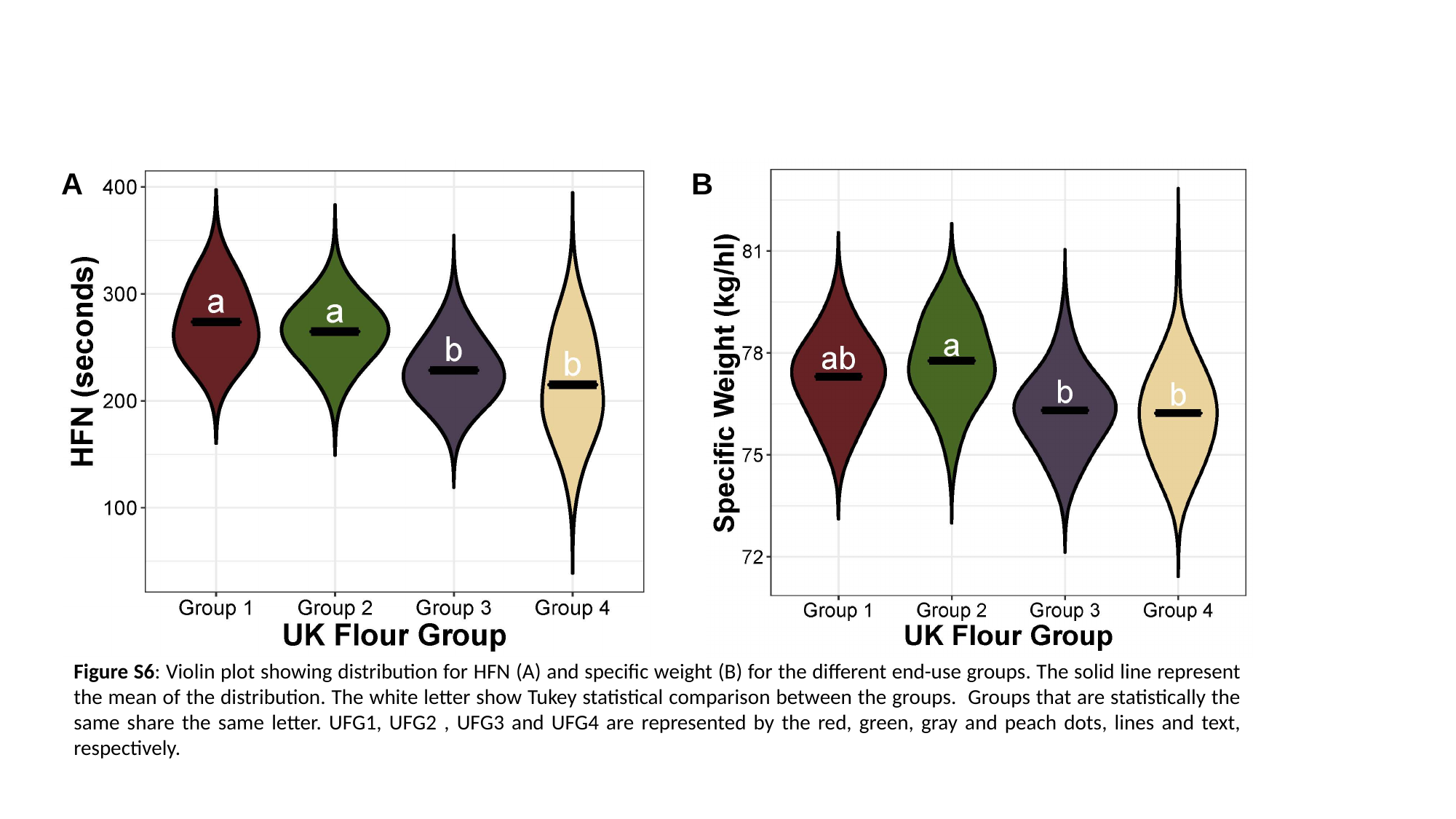

A
B
Figure S6: Violin plot showing distribution for HFN (A) and specific weight (B) for the different end-use groups. The solid line represent the mean of the distribution. The white letter show Tukey statistical comparison between the groups. Groups that are statistically the same share the same letter. UFG1, UFG2 , UFG3 and UFG4 are represented by the red, green, gray and peach dots, lines and text, respectively.

### Slide 7
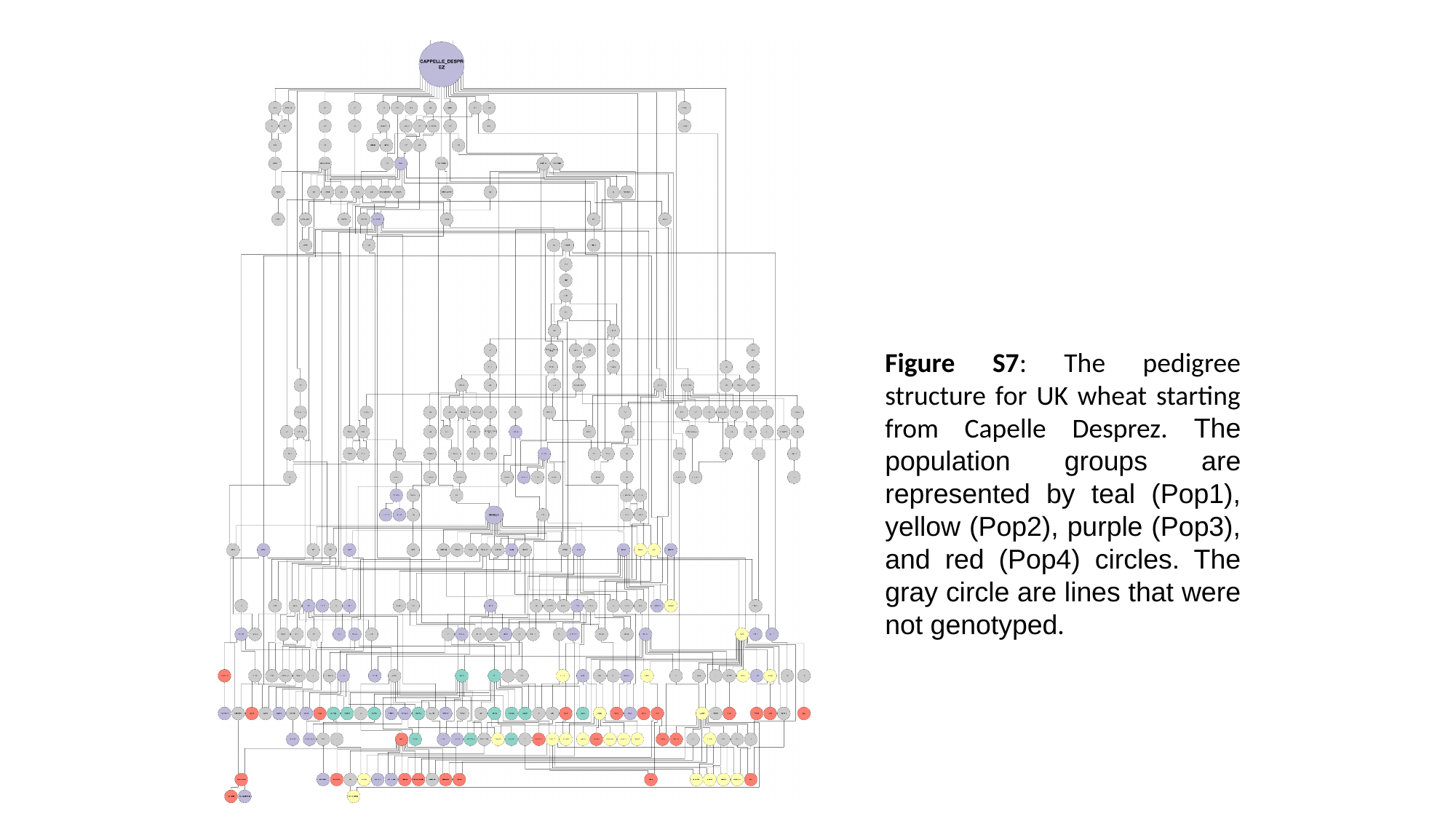

Figure S7: The pedigree structure for UK wheat starting from Capelle Desprez. The population groups are represented by teal (Pop1), yellow (Pop2), purple (Pop3), and red (Pop4) circles. The gray circle are lines that were not genotyped.
